# Tumor-derived colorectal cancer organoids induce a unique Treg cell population through direct modulation of CD4^+^ T cell differentiation

**DOI:** 10.1101/2024.07.18.604049

**Authors:** Sonia Aristin Revilla, Cynthia Lisanne Frederiks, Stefan Prekovic, Enric Mocholi, Onno Kranenburg, Paul James Coffer

## Abstract

In colorectal cancer (CRC), increased numbers of tumor-infiltrating CD4^+^ regulatory T (Treg) cells correlate with tumor development and immunotherapy failure, leading to poor prognosis. However, the molecular and cellular mechanisms governing Treg recruitment, expansion, or differentiation remain unclear. Here, we developed an *in vitro* co-culture system to assess the capacity of CRC tumors to directly modulate Treg cell differentiation. CD4^+^ T cells from Foxp3eGFP mice were co-cultured with murine tumor-derived CRC organoids, resulting in a significant increase in Treg cell numbers. This induction of Treg cells was not due to increased proliferation, but rather through differentiation of CD4^+^ T cells in a TGFβ-dependent manner. Human CRC tumor organoids similarly induced Treg cells that exhibited enhanced suppressive capacity compared to TGFβ-induced Treg cells. RNA-sequencing analysis identified distinct transcriptional profiles between CRC organoid-induced Treg cells and TGFβ-induced Treg cells, with upregulation of key functional signature genes linked to CRC Treg cells *in vivo*. High expression of genes upregulated in CRC organoid-induced Treg cells correlates with shorter progression free interval and overall survival of CRC patients, highlighting their prognostic potential. Taken together, CRC tumor organoids drive CD4^+^ differentiation to Treg cells with a phenotype resembling tumor-infiltrating Treg cells. This model can be applied to both understand the molecular mechanisms by which tumors can directly modulate CD4^+^ T cell differentiation and identify approaches to disrupt Treg cell function and stimulate anti-tumor immunity.

## Introduction

Regulatory T (Treg) cells, a specialized subset of CD4^+^ T cells, exert immunosuppressive functions crucial for maintaining immune homeostasis and self-tolerance. They are characterized by expressing FOXP3, a master transcription factor governing Treg cell differentiation and suppressive function ^1,2^. Treg cells are found in almost all peripheral tissues and adapt their transcriptional profile in response to diverse microenvironmental stresses ^3–5^. Consequently, diverse transcriptional programs related to tissue-resident Treg cell shape their role in tissue homeostasis and define their tissue-specific functions regulating multiple processes such as tissue repair and regeneration across multiple sites ^5,6^. They also exhibit cerebroprotective effects during acute experimental stroke or mitigating metabolic inflammation in adipose tissue ^5,7,8^. Through their interplay with tissue microenvironments, Treg cells ensure both immune equilibrium and tissue homeostasis.

Within the context of tumorigenesis, the accumulation of immunosuppressive Treg cells in tumor tissues poses a significant obstacle for evading immune-mediated tumor eradication ^9–11^. Notably, in solid tumors Treg cells frequently constitute 30-70% of all CD4^+^ T cells^12–14^. The increased frequency of tumor-infiltrating Treg (TI-Treg) cells correlates with unfavorable prognostic outcomes across various cancer types ^12–16^. In colorectal cancer (CRC), increased TI-Treg cell accumulation correlates with disease progression, metastasis, immunotherapy resistance, and poorer prognosis, although a causative link remains to be established ^10,11,17–20^. Despite these correlations, questions persist regarding the drivers promoting the accumulation of TI-Treg cell in tumors, TI-Treg cell mechanisms for suppressing the antitumor immune responses, and their distinguishing features compared to systemic Treg cells.

TI-Treg cell within the TME play a pivotal role in reinforcing the immunosuppressive milieu through a broad range of immune regulatory mechanisms ^9,11^. This includes inhibiting the activation and proliferation of immune cells through both contact-dependent and -independent mechanisms. They can release immunomodulatory cytokines (TGFβ, IL-10 and IL-35), cytolytic molecules (perforin and granzymes) and metabolically disruptive molecules (adenosine and cAMP) that impede effector cell function ^9,11^. Additionally, they express immune checkpoint receptors including GITR, OX-40 and LAG-3, which, through contact-dependent mechanisms, inhibit processes such as APC maturation and function ^9,11^. The reciprocal interaction between TME and Treg cells, promoting an immunosuppressive environment, elucidates their complex relationship and diverse mechanisms of immune regulation.

Several mechanisms have been proposed to explain the accumulation of Treg cells within the TME. Tumor cells release chemo-attractants that recruit Treg cells expressing specific chemokine receptors to the TME, such as the CCR5–CCL5 or CCR6–CCL20 chemokine axis ^9,20^. Different cells in the TME, such as stromal cells, can also increase the availability of the immunosuppressive cytokine TGFβ, thereby promoting the *in situ* differentiation of TI-Treg cell from CD4^+^ T cells, contributing to their accumulation within the TME ^21,22^. Moreover, the TME is characterized by nutrient depletion yet rich in metabolic by-products of cancer cells such as lactate. Lactate promotes the enrichment and regulatory function of Treg cells by modulating metabolic pathways within the TME ^11,23,24^. TI-Treg cells exhibit metabolic adaptation by enhancing FA-oxidation (FAO) and oxidative phosphorylation (OXPHOS) to sustain their survival, function, and proliferation ^11,25,26^.

Exploring CRC tumor-immune cell dynamics requires understanding the identity of TI-Treg cells and understanding the molecules and pathways driving Treg cell accumulation and function within the TME. These insights are pivotal for developing targeted therapeutic interventions to disrupt their immunosuppressive function and mitigate autoimmune toxicities associated with systemic Treg depletion ^11,27^. It is important to understand how CRC tumor-secreted factors modulate Treg cell proliferation, differentiation, and function. A molecular and mechanistic understanding of CD4^+^ T cell dynamics, especially Treg cells influenced by tumor-secreted factors, requires advanced *in vitro* models. Integrated models such as organoids and co-culture systems offer a more physiologically relevant platform to study CD4^+^ T cell responses to tumor-secreted factors ^28–32^. Despite extensive research focused on studying the behavior of tumor-reactive T cells through co-culture with tumor-organoids, investigation of Treg cell-TME dynamics *in vitro* remains limited ^31,33,34^. Here, we established a novel *in vitro* co-culture model with CRC-organoids and CD4^+^ T cells to explore how CRC tumors can directly influence CD4^+^ T cell behavior. We observed that CRC tumor-organoids can directly promote the differentiation of Treg cells. These tumor organoid-induced Treg (TO-iTreg) cells have a distinct transcriptional profile akin to *in vivo* CRC TI-Treg cells including increased expression of genes with prognostic potential in CRC. These data offer valuable insights for understanding the capacity of CRC tumors to influence CD4^+^ differentiation and the targeting of Treg cells in CRC treatment.

## Results

### CRC tumor-organoids induce CD4^+^ differentiation into Treg cells

To determine the capacity for CRC tumor-organoids to influence Treg cell fate specification *in vitro*, a co-culture model was developed utilizing CD4^+^ T cells together with CRC tumor-organoids. We evaluated the possible influence of CRC tumor-organoids in a contact-independent manner using a transwell co-culture system. Here, CD4^+^ T cells were plated in suspension alone or in co-culture with murine CRC tumor-organoids (mTO1 and mTO2) seeded in matrix droplets on the bottom of a plate (**Figure 1a**). CD4^+^ T cells were isolated from transgenic C57BL/6 Foxp3EGFP mice. Subsequently, these cells were pre-stimulated *in vitro* with anti-CD3/CD28 monoclonal antibodies and soluble IL-2, either with or without CRC tumor-organoids present. At two and five days after starting the culture, we evaluated Treg cell induction (CD4^+^ CD25^hi^ Foxp3 eGFP^+^ Treg cells) by flow cytometry (**Suppl. Figure 1a-b**). We first evaluated the influence of the matrix (BME) on the eGFP expression to explore any direct impact on Foxp3 expression. CD4^+^ T cells were cultured in transwell inserts with or without BME at the bottom of the plate. After two or five days, the control group of activated CD4^+^ T cells showed no increase in eGFP expression, regardless of the presence of BME (**Suppl. Figure. 1c**). CD4^+^ T cells cultured under these conditions were treated with TGFβ to induce Treg cells. eGFP expression was high in TGFβ-induced (TGFβ-i) Treg, independently of BME (**Suppl. Figure. 1d**). When we co-cultured CD4^+^ T cells with CRC tumor organoids, we observed a significant increase in eGFP expression compared to the control after two and five days of co-culture (**Figure 1b-c**). To measure the activation status of CD4^+^ T cells co-cultured with organoids, we assessed CD25, which is highly expressed in TI-Treg cells ^14,35^. CD25 expression was significantly higher on TO-iTreg cells compared to CD4^+^ Foxp3 eGFP^-^ T cells co-cultured with CRC tumor-organoids (**Figure 1d**).

**Figure 1.**
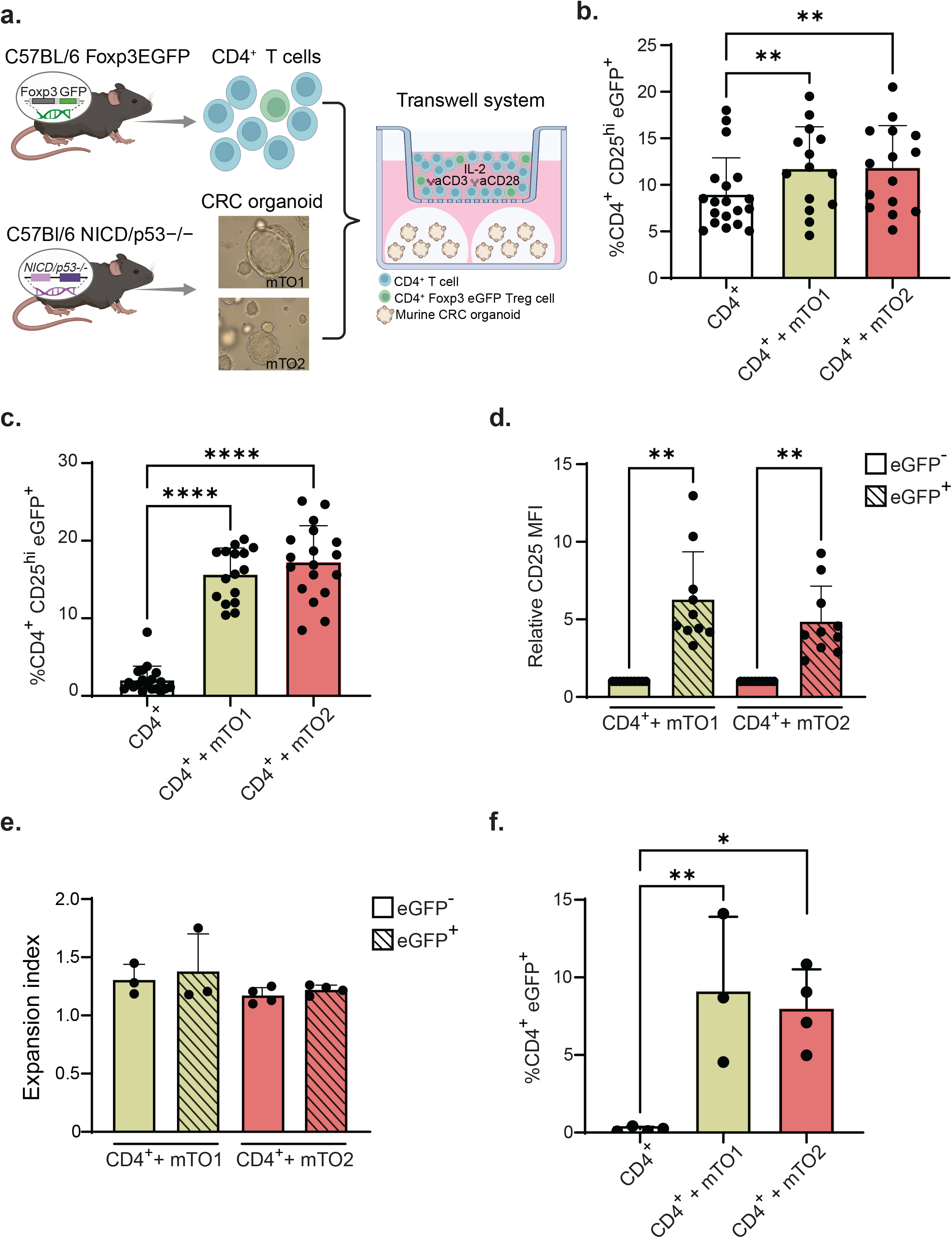
*In vitro* generation of CRC TO-iTreg cells. (**a**) Schematic illustrating the transwell co-culture system used in the experiments with CRC murine tumor organoids (mTO) and CD4^+^ T cells. (**b-c**) Percentage of CD4^+^ CD25^hi^ Foxp3 eGFP^+^ Treg cells after 2 (**b**) and 5 (**c**) days of CD4^+^ T cells culture alone or in co-culture with mTO1 and mTO2, assessed using flow cytometry. (**d**) Relative MFI of CD25 expression of eGFP^-^ (empty bars) compared to eGFP^+^ (pattern bars) CD4^+^ T cell after 5-day co-culture with mTO1 and mTO2, assessed using flow cytometry. (**e**) Expansion index of eGFP^-^ (empty bars) and eGFP^+^ (pattern bars) CD4^+^ T cells after 5-day co-culture with mTO1 and mTO2, assessed using flow cytometry. (**f**) Percentage of CD4^+^ eGFP^+^ Treg cells after 5 days of isolated CD4^+^ eGFP^-^ T cells cultured alone or in co-cultured with mTO1 and mTO2, assessed using flow cytometry. Data are represented as mean ± SD. P-values were calculated using one-way ANOVA and paired t-test analysis. *p ≤ 0.05, **p ≤ 0.01, ***p ≤ 0.001 and ****p ≤ 0.0001, ns; not significant. CD4^+^: T activated cells. mTO1 and mTO2: CRC murine tumor-organoid lines.

To evaluate whether the increased percentage of Foxp3 eGFP^+^ Treg cells was due to expansion, the proliferation of CD4^+^ T cells and Treg cells in co-culture with CRC tumor-organoids was assessed. CD4^+^ Foxp3 eGFP^-^ T cells and CD4^+^ Foxp3 eGFP^+^ Treg cells proliferated similarly (**Figure 1e**). To determine whether the induction of Foxp3 eGFP^+^ Treg cells could therefore be due to differentiation of CD4^+^ T cells, we sorted CD4^+^ Foxp3 eGFP^-^ T cells and co-cultured them with CRC tumor-organoids. We observed a significant increase in eGFP expression when CD4^+^ Foxp3 eGFP^-^ T cells were co-cultured with CRC tumor-organoids for five days compared to the control group (**Figure 1f**). Taken together, these observations demonstrate that CRC tumor-organoids can directly induce differentiation of CD4^+^ T cells to Foxp3 eGFP^+^ Treg cells in a contact-independent manner.

### CRC TO-iTreg cell generation is TGFβ-dependent and enhanced by lactate

TGFβ plays an essential role in peripheral Treg cell development and function ^22,36^. To evaluate whether there was also a role for TGFβ in the generation of TO-iTreg cells we measured TGFβ expression and release by CRC tumor-organoids cultured with or without CD4^+^ T cells. The mRNA expression levels of TGFβ in CRC tumor-organoids, cultured with or without CD4^+^ T cells, exhibited no significant difference (**Figure 2a**). To subsequently determine whether TGFβ was released from CRC tumor-organoids, we performed ELISA analysis of conditioned media collected on day five of co-culture. TGFβ was detected in conditioned media, and this was independent of the presence of activated CD4^+^ T cells (**Figure 2b**). This suggests that CRC tumor-organoids are the primary source of TGFβ secretion in this co-culture system. To further explore the relevance of these observations, we utilized TGFβ pathway inhibitors SB431542 (SB, 10μM) and LY364947 (LY, 1μM). Following a five-day incubation period, flow cytometry analysis was used to assess the percentage of Treg cells (CD4^+^ CD25^hi^ Foxp3 eGFP^+^ Treg cells). SB and LY both significantly decreased the induction of Foxp3 eGFP^+^ Treg cells in co-cultures (**Figure 2c**). To evaluate the role of lactate in promoting Treg cells enrichment, we added sodium L-lactate (10 mM) to the co-culture medium. Following a five-day incubation, we again assessed Foxp3 eGFP^+^ Treg cell induction by flow cytometry. We observed a significant increase in Treg cell induction within the co-cultures after addition of sodium L-lactate (**Figure 2d**). To determine whether lactate metabolism was essential for TO-iTreg cell generation we first utilized lactate dehydrogenase inhibitor GSK2837808A (LDHi, 10μM) which prevents the conversion of lactate to pyruvate and *vice versa* ^37^. No significant difference in Foxp3 eGFP^+^ Treg cell induction with or without LDHi was observed (**Figure 2e**). Additionally, we employed dichloroacetate (DCA, 5mM), a pyruvate dehydrogenase kinase inhibitor, to enhance pyruvate oxidation, thereby driving pyruvate into the mitochondria, reducing lactate levels ^38^. Similarly to LDHi, no significant difference in Foxp3 eGFP^+^ Treg cell induction was observed in the presence of DCA (**Figure 2f**). These findings support an essential role for TGFβ signaling in TO-iTreg cells generation. Additionally, while lactate promotes Treg cell enrichment in co-cultures, inhibiting lactate metabolism does not significantly affect Treg cell induction.

**Figure 2.**
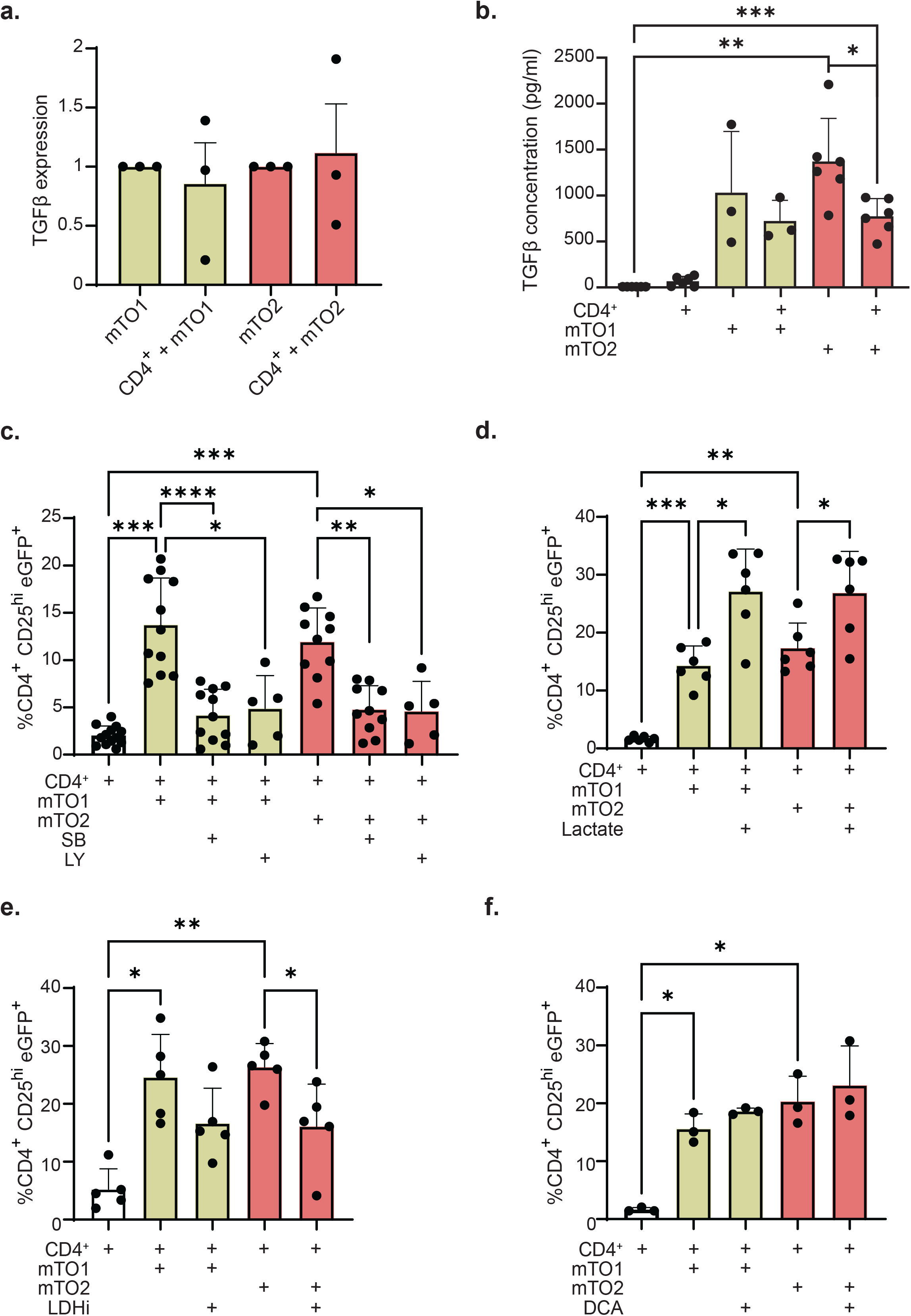
*In vitro* generation of CRC TO-iTreg cells is TGFβ dependent and enhanced by lactate. (**a**) Real-time qPCR analysis of expression levels of TGFβI, relative to B2m, in mTO1 and mTO2 organoids co-cultured with CD4^+^ T cells are shown as fold changes compared to mTO1 and mTO2 organoids cultured alone for 5 days. (**b**) TGFβ concentration in media conditioned by CD4^+^ T cells, CRC tumor organoid lines (mTO1 and mTO2), co-cultures (CD4^+^ + mTO1 and mTO2), and identically treated control medium with BME after a 5-day culture, was assessed by ELISA. (**c-f**) Percentage of CD4^+^ CD25^hi^ Foxp3 eGFP^+^ Treg cells after 5 days of CD4^+^ T cells culture alone or in co-culture with mTO1 and mTO2, supplemented with (**c**) TGFβI receptor kinase inhibitors, SB-431542 (SB, 10μM) and LY364947 (LY, 1μM), (**d**) sodium L-lactate (Lactate, 10mM), (**e**) LHD inhibitor GSK2837808A (LDHi, 10μM), and (**f**) dichloroacetate (DCA, 5mM), assessed using flow cytometry. Data are represented as mean ± SD. P-values were calculated using one-way ANOVA. *p ≤ 0.05, **p ≤ 0.01, ***p ≤ 0.001 and ****p ≤ 0.0001, ns; not significant. CD4^+^: T activated cells. mTO1 and mTO2: CRC murine tumor-organoid lines.

### Human CRC TO-iTreg cells have enhanced immunosuppressive potential *in vitro*

To extend and validate our observations, we developed a transwell co-culture system utilizing human CD4^+^ T cells derived from human cord blood. After five-days of co-culture, we assessed the proportion of Treg cells (CD4^+^ FOXP3^+^ Treg cells) by flow cytometry. Similarly to the murine system, the expression of FOXP3 in CD4^+^ CD25^+^ T cells was significantly increased in the presence of CRC tumor-organoids compared to the control group of activated CD4^+^ T cells (**Figure 3a**). CRC tumor-organoids cultured with CD4^+^ T cells expressed lower levels of TGFβ compared to those cultured alone (**Figure 3b**). The addition of SB or LY again led to reduced Treg cell induction in the co-cultures with both CRC tumor-organoids as observed in the murine model (**Figure 3c**). However, in contrast to our observations in the murine model, lactate supplementation did not result in an increased proportion of CD4^+^ FOXP3^+^ Treg cells (**Suppl. Figure. 2a**) and inhibiting LDH had minor impact on the CD4^+^ FOXP3^+^ Treg cell percentage in any of the co-culture conditions (**Suppl. Figure. 2b**). To compare the immunosuppressive functions of hTO- and TGFβ-induced Treg cells, we performed an *in vitro* suppression assay ^39^. Here, we cultured CTV-stained PBMCs with different ratios of hTO-iTreg cells or TGFβ iTreg cells, either at 30% or 80% of the total CD4^+^ T cells. The proliferation of the PBMCs was quantified by flow cytometry as described earlier. A reduction in PBMC proliferation was directly proportional to the increasing number of TGFβ- or hTO-iTreg cells (**Suppl. Figure. 2c**). PBMCs co-cultured with hTO-iTreg cells demonstrated a significant reduction in proliferation compared to PBMCs alone (**Figure 3d-e**). Additionally, hTO-iTreg cells co-cultured with PBMCs showed a stronger suppressive effect compared to those co-cultured with an equal number of TGFβ-iTreg cells (**Figure 3d-e**).

**Figure 3.**
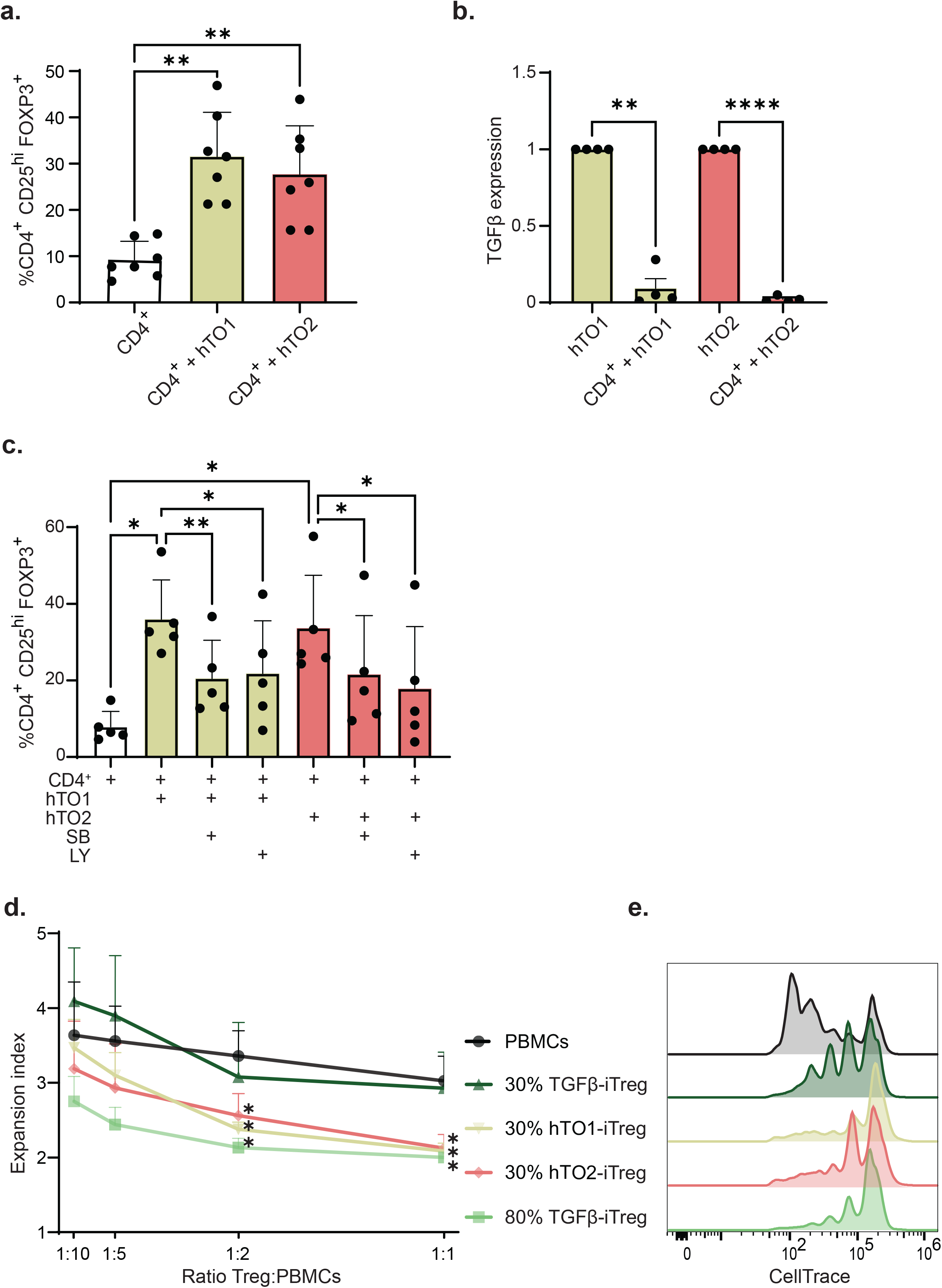
Human CRC tumors-organoids directly induce immunosuppressive Treg cells. (**a**) Percentage of CD4^+^ CD25^+^ FOXP3^+^ Treg cells after 5 days of cord blood isolated CD4^+^ T cells culture alone or in co-culture with human CRC tumor organoid lines (hTO1 and hTO2), assessed using flow cytometry. (**b**) Real-time qPCR analysis of expression levels of TGFβI, relative to B2m, in hTO1 and hTO2 organoids co-cultured with CD4^+^ T cells are shown as fold changes compared to hTO1 and hTO2 organoids cultured alone for 5 days. (**d**) Percentage of CD4^+^ CD25^+^ FOXP3^+^ Treg cells after 5 days of CD4^+^ T cells culture alone or in co-culture with hTO1 and hTO2, supplemented with TGFβI receptor kinase inhibitors, SB-431542 (SB, 10μM) and LY364947 (LY, 1μM), assessed using flow cytometry. (**d**) Survival curve showing the expansion index of different ratios of CTV-stained PBMCs cultured alone or co-cultured with different ratios of diluted 30% TGFβ iTreg cells, 30% hTO1-iTreg cells, 30% hTO2-iTreg cells and 80% TGFβ iTreg cells after 4-day co-culture, assessed using flow cytometry (n = 3, different donors). (**e**) Representative histograms depicting the proliferation of CTV-stained PBMCs alone or with diluted 30% TGFβ iTreg cells, 30% hTO1-iTreg cells, 30% hTO2-iTreg cells and 80% TGFβ iTreg cells. Data are represented as mean ± SD. P-values were calculated using one-way ANOVA. *p ≤ 0.05, **p ≤ 0.01, ***p ≤ 0.001 and ****p ≤ 0.0001, ns; not significant. CD4^+^: T activated cells, hTO1 and hTO2: CRC human tumor-organoid lines.

### CRC TO-iTreg cells express Treg cell associated markers

To define the transcriptional identity of *in vitro* mTO-iTreg cells we performed bulk RNA-sequencing. To this end, we employed a staged cell sorting approach at different time points. Firstly, Foxp3 eGFP^-^ CD4^+^ T cells were isolated on day 0, prior to co-culture establishment; secondly, CD4^+^ Foxp3 eGFP^+^ TGFβ-iTreg cells were isolated on day 5 of culture; and thirdly, CD4^+^ Foxp3 eGFP^+^ mTO-iTreg cells were isolated from the co-culture on day 5. Principle Component Analysis (PCA) identified the mTO-iTreg cell group as a distinct cluster, separate from TGFβ iTreg cells (**Figure 4a**). To assess the expression of core Treg cell signature genes among the three CD4^+^ T cell populations, a gene list was generated informed by existing literature ^11,40,41^. Both mTO-iTreg cells and TGFβ-iTreg cells exhibit significantly elevated expression levels of core Treg cell signature genes when compared to the control group of eGFP^-^ CD4^+^ T cells (**Figure 4b-c**). *Foxp3* and *IL2RA* (CD25), key markers for identifying and characterizing Treg cells, show higher expression levels in TI-Treg cells in comparison to Treg cells present in the peripheral systemic circulation ^14,35,42^. *Foxp3* exhibited significantly higher expression in mTO-iTreg cells compared to TGFβ-iTreg cells (**Figure 4d**). While *IL2RA* displayed significantly elevated expression in both Treg cell populations relative to control (**Figure 4e**). To identify similarities and differences between TGFβ-iTreg cells and mTO-iTreg cells, we analyzed differentially expressed genes (DEGs; **Figure 4f**). We identified 126 genes exclusive to TGFβ-iTreg cells, while 1403 genes exhibited differential expression specifically in mTO-iTreg cells compared to the control (**Figure 4f**). An additional 126 genes were expressed in both mTO-iTreg cells and TGFβ-iTreg cells compared to the control group (**Suppl. Figure 3a-b**). Analysis of the 126 genes shared between mTO-iTreg cells and TGFβ-iTreg cells by Gene Ontology (GO) term analysis identified genes significantly contributing to lymphocyte activation, differentiation, and cytokine response pathways (**Suppl. Figure 3c**). Additionally, enriched GO terms linked these genes to the regulation of catabolic processes, suggesting involvement in modulating essential metabolic pathways crucial for Treg cell maintenance and function (**Suppl. Figure 3c**). Taken together, mTO-iTreg cells and TGFβ-iTreg cells are transcriptionally distinct.

**Figure 4.**
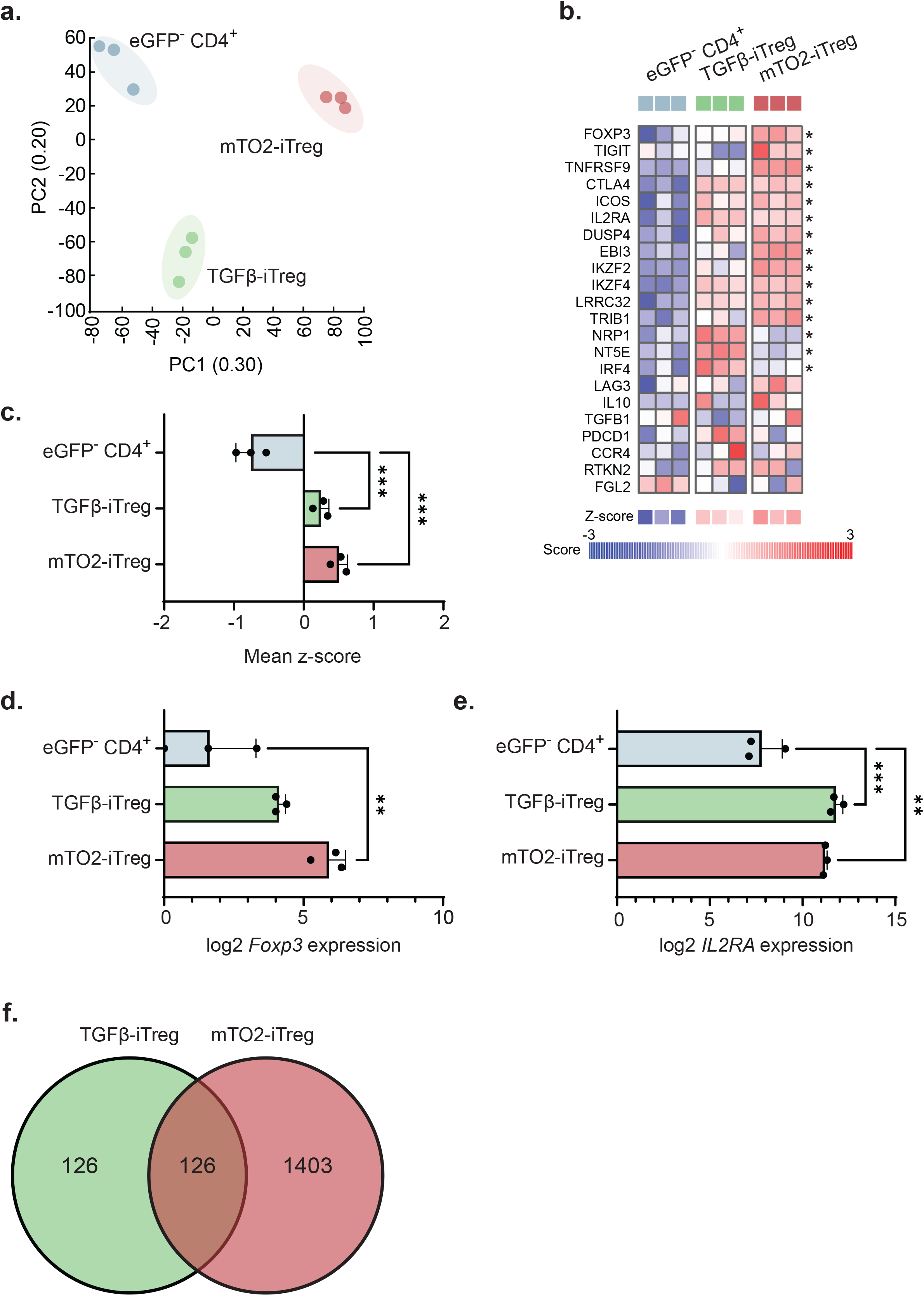
*In vitro* CRC TO-iTreg cells express Treg cells associated markers. (**a**) PCA performed on splenic eGFP^-^ CD4^+^ T cells (blue), TGFβ-iTreg cells (green) and mTO2-iTreg cells (red). (**b**) Heatmap and (**c**) bar chart quantifying (mean ± SD) the signature z-score of core Treg cells signature genes expression in splenic eGFP^-^ CD4^+^ T cells (blue), TGFβ-iTreg cells (green) and mTO2-iTreg cells (red). (**d-e**) Bar chart represent the log2-fold change (mean ± SD) in (**d**) *Foxp3* and (**e**) *IL2RA* gene expression between splenic eGFP^-^ CD4^+^ T cells (blue), TGFβ-iTreg cells (green) and mTO2-iTreg cells (red). (**f**) Venn diagram analysis of DEGs between control splenic eGFP^-^ CD4^+^ T cells (control) and TGFβ-iTreg cells (green) or mTO2-iTreg cells (red) (ANOVA, FDR, p ≤ 0.05). P-values were calculated using one-way ANOVA. *p ≤ 0.05, **p ≤ 0.01, ***p ≤ 0.001 and ****p ≤ 0.0001, ns; not significant. mTO2: CRC murine tumor-organoid line.

### CRC TO-iTreg cells upregulate signature genes associated with tumor-infiltrating Treg cells *in vivo*

To assess the differences between *in vitro* mTO-iTreg cells and TGFβ-iTreg cells, we conducted a differential gene expression analysis. A pool of 1155 significantly (p≤0.05) DEGs were identified with differential expression between mTO-iTreg cells and TGFβ-iTreg cells (**Figure 5a**). From this pool, we extracted the 56 most significantly DEGs (p≤0.01) **(Suppl. Figure 4a**), including *Tgm2, Pttg1*, and *Fibp*, which have been associated with TI-Treg cells in the TME of other cancers ^43–47^. To delineate the biological processes associated with the 1155 DEGs, we conducted a pathway analysis (**Suppl. Figure 4b**). This analysis revealed differential expression of genes involved in biosynthetic processes, such as translation or peptides biosynthesis; cellular regulation and activation, such as lymphocyte activation or regulation of immune system process; cellular organization and transport, including protein transport or leukocyte cell-cell adhesion; and, metabolic processes, such as canonical glycolysis or oxidative phosphorylation (**Suppl. Figure 4b**). mTO-iTreg cells exhibit upregulated expression of glycolysis pathway genes within this GO term annotation list (**Suppl. Figure 4c-d**). Additionally, our analysis of cellular regulation and activation pathways revealed differences in transcription factors (TFs) associated with Treg cells between TGFβ-Treg and mTO-Treg cells. mTO-iTreg cells show significant increased expression of genes including *Foxp3* (FOXP3), *Eomes* (EOMES), *Hic1* (HIC1), *Gata3* (GATA3), *Xbp1* (XBP1) and *Ikzf2* (Helios), alongside significant downregulation of transcription factors such as *Irf4* (IRF4), *Rela* (RELA), *Maf* (C-MAF), *Rorc* (RORγt), and *Foxo1* (FOXO1), which are crucial for promoting differentiation and modulating immune responses (**Figure 5b** and **Suppl. Figure 5e**) ^48–50^. We subsequently sought to determine whether mTO-iTreg cells exhibit closer transcriptional similarities with *in vivo* CRC TI-Treg cells compared to TGFβ-iTreg cells. To this end, we investigated the expression of genes previously reported to be more highly expressed in *in vivo* CRC TI-Treg cells as compared to peripheral Treg cells in healthy tissue or peripheral blood ^10,11^. The majority of these genes were upregulated in mTO-iTreg cells when compared to TGFβ-iTreg cells (**Figure c** and **Suppl. Figure 5f**). mTO-iTreg cells exhibit upregulation of functional signature genes associated with activated and suppressive TI-Treg cells, including *Tnfrsf4* (OX40), *Tnfrsf18* (GITR), *Tnfrsf9* (4-1BB), and *Tigit* (TIGIT), in comparison to TGFβ-iTreg cells (**Figure 5c**) ^40,41^. The increased expression of some of these upregulated genes was verified at the protein level by flow cytometry analysis. OX40, GITR, FAS, and CD27 protein levels were higher in mTO-iTreg cells compared to TGFβ-iTreg cells, validating the observed transcriptional increases (**Figure 5d**). To investigate whether there was a potential association between upregulated genes in mTO-iTreg cells and CRC prognosis, we identified the top 141 most upregulated genes (with >3 log-fold change) in mTO-iTreg cells (**Suppl. Figure. 5**). Subsequently, using the gene expression data from the TCGA of CRC tumors we calculated the mean z-score per patient for the abovementioned upregulated genes. This cohort was stratified into low or high expression of the selected upregulated genes in mTO-iTreg cells. Kaplan–Meier analysis, which compares survival differences between the two groups of CRC patients, demonstrated a significant correlation between high expression of these selected genes and a shortened progression-free interval (PFI) (**Figure 5e**) and overall survival (OS) (**Figure 5f**) in CRC tumors. Overall, these observations demonstrate a unique transcriptional profile of mTO-iTreg cells with increased similarity to *in vivo* CRC TI-Treg cells compared to TGFβ-iTreg cells, indicating their potential to inform CRC prognosis.

**Figure 5.**
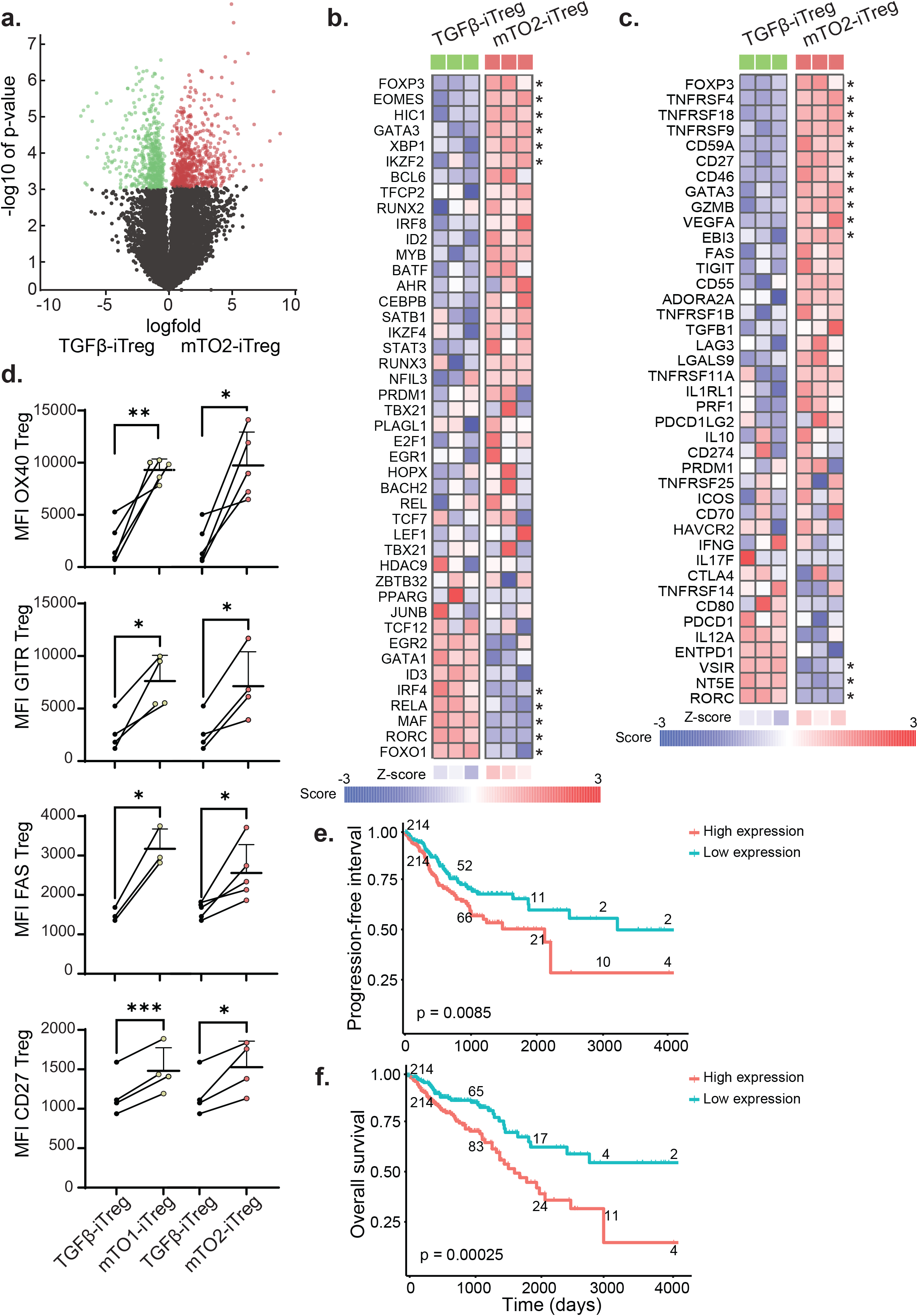
*In vitro* CRC TO-iTreg cells upregulate functional signature genes associated with CRC TI-Treg cells *in vivo*. (**a**) Volcano plot displays gene expression fold change against statistical significance, highlighting significant upregulation in mTO2-iTreg cells (red) compared TGFβ-iTreg cells (green) (ANOVA, FDR, p ≤ 0.05). (**b**) Heatmap of the expression of TFs genes associated with Tregs cells in mTO2-iTreg cells compared to TGFβ-iTreg cells. (**c**) Heatmap of the expression of genes associated with *in vivo* CRC TI-Treg cells in mTO2-iTreg cells compared to TGFβ-iTreg cells. (**d**) MFI of OX40, GITR, FAS, and CD27 expression in mTO1- and mTO2-iTreg cells compared to TGFβ-iTreg cells. (**e-f**) Kaplan-Meier curves for CRC patients of the TCGA cohort, using (**e**) progression-free interval and (**f**) overall survival with patients split based on the z-score of the identified signature into two groups (low and high gene expression). The numbers included in the curves indicate the number of CRC patients per time point. P-values were calculated using one-way ANOVA and paired t-test analysis. *p ≤ 0.05, **p ≤ 0.01, ***p ≤ 0.001, and ****p ≤ 0.0001, ns; not significant. mTO1 and mTO2: CRC murine tumor-organoid lines.

## Discussion

*In vivo*, Treg cell differentiation and accumulation within the CRC TME significantly hinders immune surveillance, ultimately promoting tumor progression ^11,20^.However, the precise mechanisms by which CRC directly influences Treg cell phenotype and function remain largely unknown. Insight into TI-Treg cell identity, mechanisms of accumulation, and function in the TME is critical for targeting them in immunotherapy. Here, we present a novel *in vitro* co-culture model designed to identify and assess the direct effects of CRC-derived factors on CD4^+^ T cell homeostasis, particularly in modulating Treg cell dynamics. CRC-organoids were found to directly increase the number of TO-iTreg cells through *de novo* differentiation of CD4^+^ T cells in a cell contact-independent and TGFβ-dependent manner. Moreover, *in vitro* generated TO-iTreg cells are transcriptionally and phenotypically distinct when compared to TGFβ-iTreg cells. This includes the expression of critical regulatory genes and distinct cell surface marker profiles, thereby making TO-iTreg cells more closely resembling TI-Treg cells found within the TME. High expression of some of these TO-iTreg genes in CRC tumors correlates with shorter progression-free interval (PFI) and overall survival (OS), emphasizing their prognostic significance.

Recent studies highlight the relevance of the CRC tumor organoid-immune cell co-culture models, which provide a closer approximation to the *in vivo* TME and advancing therapeutic interventions ^31,33,34^. While significant attention has been given to studying tumor-reactive T cells using a co-culture system with CRC tumor-organoids, our study is the first utilizing a co-culture system evaluating the capacity of tumor organoids to directly modulate Treg cell differentiation. This model holds potential to enhance our understanding of how CRC tumors can directly modulate CD4^+^ T cells homeostasis resulting in increased Treg cell differentiation. Furthermore, it offers a valuable *in vitro* platform for identifying biomarkers associated with Treg cells in CRC, potentially allowing *in vitro* screening strategies to identifying and target tumor-specific Treg cells.

Using a contact-independent *in vitro* co-culture system, we confirmed the capacity of CRC tumor-organoids to induce TO-iTreg cells in a contact-independent manner through conversion of CD4^+^ T cells to Treg cells. Here, the release of TGFβ by CRC tumor-organoids plays a central role (**Figures 2b, 3c**). Tauriello *et al*. demonstrated the regulatory capacity of TGFβ in modulating T cell behavior and its implications in CRC tumor metastasis utilizing a murine model of metastatic CRC ^51^. The resultant metastatic intestinal tumors exhibited elevated TGFβ activation leading to T cell exclusion and suppressed effector T cell response. In this study, CAFs were identified as the primary producers of TGFβ ^51^. However, here we found that CRC tumors can themselves directly regulate Treg cell differentiation through the production of TGFβ in the absence of any stromal support. In line with our results Yamada *et al*. demonstrated that CRC tumor cell-derived extracellular vesicles, rich in TGFβ1, suppressed Jurkat cells proliferation and induced a Treg-like cell phenotype characterized by the upregulation of genes including *FoxP3, CTLA-4, LAG3, IL-10, PRF1*, and *GZMB* ^52^. Lactate, a by-product of cancer cell glycolysis released into the TME, hinders the aerobic glycolysis-dependent activation and proliferation of murine and human effector T cells, thereby inhibiting tumor immunosurveillance and promoting tumor growth ^24,53^. In contrast, lactate facilitates Treg cell differentiation, sustaining their suppressive function through metabolic reprogramming, as primarily demonstrated in studies on murine Treg cells ^23,24^. In our study, lactate significantly enhances the induction of mTO-Treg cells but not hTO-Treg cells (**Figures 2d, Suppl. Figure. 2a**). Previous research has linked high lactate concentrations with increased suppressive capacity in human Treg cells ^54^. However, mechanisms governing human Treg cell differentiation and function in various metabolic environments are still largely unknown. Understanding the distinct origins and energy requirements of Treg cells and effector T cells in the TME may help identify therapeutic targets. For instance, TGFβ inhibition with other therapies holds promise for CRC treatment by dampening TI-Treg cells and bolstering effector T cell responses, as seen in other cancers ^55^.

TGFβ signaling is associated with stable *Foxp3* expression, and therefore Treg cell differentiation and function ^22,36^. Our results confirm the critical role of TGFβ in promoting *Foxp3* expression in TO-iTreg cells. Nevertheless, RNA-seq analysis demonstrates a distinct transcriptional phenotype between mTO-iTreg cells and TGFβ-iTreg cells, suggesting that while mTO-iTreg cells require TGFβ signaling (**Figure 2c**), there must be additional contributing factors driving the separation of mTO-iTreg cells from TGFβ iTreg cells at the RNA level (**Figure 4a**). Moreover, mTO-iTreg cells exhibit a greater similarity to *in vivo* CRC TI-Treg cells. The increased expression of genes associated with immune activation and regulatory functions such as *Cd2* (CD2), *GZMB* (Granzyme B), *Cd46* (CD46), *Cd59a* (CD59), *Cd27* (CD27), *Tnfrsf4* (OX40), *Tnfrsf18* (GITR), *Tnfrsf9* (4-1BB), and *Tigit* (TIGIT), indicates increased activation and suppressive functionality in mTO-iTreg cells ^40,41^. This is supported by the differential expression of TFs observed in mTO-iTreg cells (**Figure. 5b**). Key factors *Gata3* (GATA3) and *Ikzf2* (Helios) are critical for maintaining Foxp3 expression and lineage stability ^56,57^. Decreased Foxo1 facilitates Treg cells migration to non-lymphoid organs, while Gata3^+^ Helios^+^ Treg cells, constituting approximately one-third of colonic Treg cells, are pivotal in fate determination and accumulating in inflamed tissues, including CRC ^56–58^. Additionally, increased levels of *Helios, Eomes, Hic1* and *Bcl6* along with decreased *Foxo1*, contribute to Treg effector suppressive capabilities impairing antitumor immune responses ^57–61^. The expression of *Hic1* (HIC1), *Foxp3* (FOXP3) and *Tcf7* (TCF-1), together with the downregulation of *Maf* (MAF), *Rorc* (RORγt), *Il17* (IL-17), and *Ifng* (IFNγ) (**Figure 5b-c**) —all crucial in regulating intestinal immune homeostasis— suggests the suppression of proinflammatory Th17-like differentiation and absence of β-catenin pathway activation in mTO-iTreg cells ^62–65^. Differentially expressed mTO-iTreg cell genes also include those associated with metabolic processes such as glycolysis and oxidative phosphorylation, potentially contributing to the energy requirements and metabolic adaptations of mTO-iTreg cells within the *in vitro* TME (**Suppl. Figure. 4b-d**). Additionally, involvement in biosynthetic processes such as translation and peptide biosynthesis suggest potential roles in producing regulatory molecules and effector proteins crucial for mTO-iTreg cells function within the *in vitro* TME. Aligned with these findings, De Ponte Conti *et al*. demonstrated that within distinct TILs subtypes, TI-Treg cells and tumor-reactive CD8^+^ T cells exhibit an increased translational activity upon activation^66^. Subsequently, TI-Treg cells upregulated genes associated with their proliferative capacity and immunosuppressive phenotype ^66^. In conclusion, the transcriptional changes in mTO-iTreg cells indicate a highly suppressive and stable phenotype, resembling tissue-resident and effector Treg cells found in CRC tumors. This suggests their contribution to the establishment and maintenance of an immunosuppressive TME *in vitro*. Understanding the biological relevance of differentially expressed genes in TO-iTreg cells may provide novel insights into how they exert immunosuppressive functions and influence tumor progression within the TME, offering unique therapeutic targets for intervention.

Elevated levels of immunosuppressive TI-Treg cells within CRC tumors are associated with poorer prognosis in patients ^11,17–20^. Within the pool of significantly upregulated genes in mTO-iTreg cells, *TGM2, PTTG1* and *FIBP* have previously been associated with TI-Treg cells in the TME of other cancers. High *TGM2* expression in gastric or pancreatic cancer is associated to increased Treg cells presence and a worse prognosis ^43,44^. *PTTG1* expression is found in immunosuppressive Treg cells in triple-negative breast cancer and associated with significant Treg cell infiltration in liver tumors ^45,46^. Additionally, high expression of FIBP correlates with elevated frequency of Treg cells in acute myeloid leukemia, serving as a prognostic biomarker indicating a poor prognosis ^47^. In line with these observations, our study showed that elevated expression of the most upregulated genes in mTO-iTreg cells correlates with unfavorable disease outcomes for CRC patients (**Figure 5e-f**). Correlation of these genes with poor prognosis in CRC underscores their potential as prognostic markers and emphasizes the relevance of the *in vitro* model. This model provides a controlled platform to explore how these genes influence Treg cell behavior and impact disease progression. This knowledge is essential for developing targeted therapies that manipulate immune responses in CRC, potentially leading to the development of personalized treatment strategies aimed at improving outcomes for CRC patients.

Taken together, we present a novel *in vitro* approach to generate and analyze CRC tumor-associated Treg cells. This enables investigations into CRC-immune cell dynamics and targeted strategies crucial for exploring T cell-based therapies to disrupt Treg cell function and enhance effector T cell anti-tumor responses within the TME. Ultimately, this may help drive precision medicine in CRC treatment and improve patient outcomes.

## Acknowledgments

The authors thank Liza Wijler for providing the mouse tumor organoids, and Maria Lamprou, Danai Kokkinidi, and Simona Golemanova for their experimental assistance. We also thank all members of the Coffer, Kranenburg, Prekovic and Westendorp groups for helpful discussions, the Hubrecht Institute FACS facility, Single Cell Discoveries B.V., and the R2 platform for their support and GDL Utrecht employees for assistance with the mice. All illustrations were created with BioRender.com. This work was supported in part by a Worldwide Cancer Research grant (Reference: 19-0371). The funding agencies played no role in the design, reviewing, or writing of the manuscript.

## Author contributions

S.A.R. designed, performed, analyzed experiments, prepared figures, and wrote the manuscript. C.L.F. performed and analyzed experiments. S.P. assisted with RNA sequencing analysis. E.M. and O.K. assisted with experimental design and supervised the research. P.J.C. supervised the research, helped in experimental design, and wrote the manuscript. All authors have read and agreed to the published version of the manuscript.

## Material and methods

### Murine CD4^+^ T cell isolation

Transgenic B6-Foxp3^EGFP^ mice (B6.Cg-*Foxp3*^*tm2(EGFP)Tch*^/J, The Jackson Laboratory, stock no. 006772) were euthanized for the isolation of CD4^+^ T cells. These mice express EGFP under the same promoter of the Treg cell-specific transcription factor Foxp3 (Foxp3 eGFP). Thus, EGFP can be used as a surrogate marker of Treg cells presence, which enables quick assessment of Treg cell numbers by flow cytometry analysis ^67^. From each mouse, lymph nodes (inguinal, brachial, axillary, and cervical) and spleen were harvested individually and smashed against the mesh (70 μm) of a sterile cell strainer in a cell-culture dish containing ice-cold MACs buffer (2% heat-inactivated FBS (Gemini Bio-Products), 2mM EDTA in PBS). The total number of cells isolated was assessed using a TC20 automated cell counter (Bio-Rad) using Trypan Blue dye. CD4^+^ T cells were positively isolated with mouse CD4 (L3T4) microbeads (Miltenyi Biotec) following the manufacture’s protocol. For each mouse, one LC column (Miltenyi Biotec) was placed tightly in the QuadroMACS™ Separator (Miltenyi Biotec). Total alive CD4^+^ T cell number was determined by TC20 automated cell counter using Trypan Blue dye.

### Human CD4^+^ T cell isolation

CD4^+^ T cells were isolated from cord blood of healthy donors obtained according to the Declaration of Helsinki. The collection protocol was approved by the Ethical Committee of the Utrecht Medical Center (UMCU) in the Netherlands. After Ficoll-Paque (GE Healthcare) gradient separation, cord blood mononuclear cells (CBMCs) were cryopreserved for later use. CD4^+^ T cells were isolated from the CBMCs fraction in MACs buffer using the MagniSort human CD4^+^ T cell enrichment kit (Thermo Fisher) and BD IMag Cell Separation Magnet (BD Biosciences) following the manufacture’s protocol. Total alive CD4+ T cell number was determined by TC20 automated cell counter using Trypan Blue dye.

### CRC tumor-organoid culture

CRC murine tumor-organoid (mTO) lines were derived from spontaneous colon tumors originated in a transgenic mouse model with conditional activation of the Notch1 receptor and deletion of p53 in the digestive epithelium (NICD/p53^-/-^) ^68,69^. Exome sequencing revealed mutations in either the *Ctnnb1* or *Apc* genes, indicating classical Wnt pathway activation ^68^. CRC human tumor-organoid (hTO) lines used for this study were established and characterized, as previously described ^70,71^. The patient’s primary CRC tumor samples were obtained during a colon resection for primary adenocarcinoma within the biobanking protocol HUB-Cancer TcBio#12-09, which was approved by the medical ethical committee of the UMCU in the Netherlands. Written informed consent was obtained from all patients. See Table 1 for an overview of all hTOs used in this study including their clinical parameters and mutational status.

**Table 1.** Clinical and mutational characteristics of the patient-derived CRC hTOs used in this study.

| Organoid ID | Named | CRC Stage | CRC Location | Gender | Age | Origin | APC | P53 | Kras | BRAF | SMAD4 | RNF43 | ARID1A | MSI-status |
| --- | --- | --- | --- | --- | --- | --- | --- | --- | --- | --- | --- | --- | --- | --- |
| Tor10 | hTO1 | T4N2M1 | Rectum | Female | 51 | Primary |  | S215I | G12A | V600E | F339L | P441Afs | Q473* | MMR+ |
| Tor18 | hTO2 | Unknown | Caecum | Male | 70 | Primary | L1489Y | I255N |  |  |  |  |  | MMR+ |

Murine and human CRC tumor-organoids were passaged once a week and their corresponding CRC tumor-organoid medium was refreshed every three days. To passage CRC tumor-organoids, BME (R&D systems) was dispersed by pipetting and washed with pre-cold PBS. CRC tumor-organoids were resuspended thoroughly with pre-warmed TrypLE-Express (Gibco), incubated for 5 minutes at 37°C and mechanically sheared to obtain a single cell suspension. Cells were then washed and resuspended in 50% BME and 50% medium to the right ratio for passaging (mTOs: 1:15 and 1:25. hTOs: 1:8 and 1:12). Cells were plated as droplets in pre-warmed culture plate and incubated upside down for approximately 30 minutes at 37°C and 5% CO_2_. After matrix solidification, cells were overlaid with their corresponding CRC tumor-organoid medium. The basal BM2 medium for murine and human CRC tumor-organoids culture consisted of Advanced Dulbecco’s Modified Eagle Medium (DMEM)/F12 medium (Gibco) supplemented with 10 mM N’-2-Hydroxyethylpiperazine-N’-2 ethanesulphonic acid (HEPES, Lonza), 2 mM Glutamax (Gibco), 50 U/ml penicillin-streptomycin (Gibco), 100 ng/ml Noggin conditioned medium (produced by lentiviral transfection), 1 mM N-acetylcysteine (Sigma) and 2% B27 serum free supplement (Gibco). Then, BM2 mouse (BM2m) medium was further enriched with 10 nM murine recombinant fibroblast growing factor (PeproTech) while BM2 human (BM2h) medium included 500 nM A8301 (MedChemExpress) and 10 μM SB202190 (ApexBio).

### Co-culture of CD4^+^ T cells and CRC-organoids

To establish the co-cultures (**Figure 1a**), first, two days old CRC tumor-organoids were retrieved from the BME droplets by adding dispase II (1 mg/ml, Gibco) directly to the medium, followed by a 10-minute incubate at 37°C with 5% CO_2_. Then, the collected CRC tumor-organoids were washed with pre-cold PBS and counted. CRC tumor-organoid were then embedded in a mixture of 50% BME and 50% BM2^-^ medium in pre-warmed plates (Corning). After placement, the plates were incubated upside down for 30 minutes at 37°C to solidify BME. Following this, CRC tumor-organoid cultured with or without CD4^+^ T cells were covered with the corresponding T activation media, BM2m or BM2h culture media supplemented with soluble IL-2 (20 U/ml, PeproTech) and functional grade anti-CD3 (aCD3, 1 μg/mL) and anti-CD28 (aCD28, 1 μg/mL) murine or human monoclonal antibodies (Invitrogen). Subsequently, the co-cultures were established at a ratio of 1:2,5 CRC tumor-organoids to CD4^+^ T cells. CD4^+^ T cells were isolated following established protocols, and their viability was determined prior to co-culture initiation. In instances of co-culture conditions, CD4^+^ T cells were suspended in T activation medium and places in hanging inserts (Corning) within the well plate for culture initiation. In control conditions where CD4^+^ T cells were cultured without CRC tumor-organoids, CD4^+^ T cells were suspended in either T activation medium or Treg induction medium (BM2m or BM2h culture media supplemented with 10 ng/ml recombinant TGFβ (R&B systems), soluble IL-2 (300 U/ml, PeproTech) and functional grade anti-CD3 (aCD3, 1 μg/mL) and anti-CD28 (aCD28, 1 μg/mL) murine or human monoclonal antibodies (Invitrogen)). These conditions were also seeded in hanging inserts (Corning) within the well plate for culture initiation. All conditions were incubated at 37°C for a period of 5 days. Following the 5-day incubation period, medium and/or cells were harvested for further experiments.

The various culture media were enriched with the following supplements: kinase inhibitors targeting the TGFβ pathway, including SB431542 (SB, 10 μM) (STEMCELL technologies) and LY364947 (LY, 1 μM) (Sigma-Aldrich); sodium L-Lactate (Lactate, 10 mM) (Sigma-Aldrich); a lactate dehydrogenase inhibitor, GSK 2837808A (LDHi, 10 μM) (Tocris Bioscience); or dichloroacetate (DCA, 5 mM) (Tocris Bioscience).

### qRT-PCR

Five days old CRC tumor-organoids, either cultured alone or with CD4^+^ T cells, were retrieved from the BME droplets and washed with pre-cold PBS. The pellet was then resuspended in 350ul of RLT lysis buffer provided in the RNeasy Mini Kit (Quiagen). RNA was isolated using the RNeasy Mini Kit following the manufacturer’s instructions. RNA quantification was conducted using the Qubit Flurometer 3.0 (Invitrogen) also following the manufacturer’s specifications. Long-term storage of RNA samples was at -80°C. 1ug of total RNA was used to synthesized cDNA using iScript cDNA synthesis kit (Bio-Rad) accordingly to the manufacturer’s protocol. cDNA samples were stored at -20°C. Real-time quantitative PCR (qPCR) was performed using forward and reverse primers (10uM, Integrated DNA Technologies), which sequences are specified below, to amplify TGFβI. Levels of gene expression were quantified using SYBR® Green qPCR master mix method (Thermo Fisher Scientific) in a light-based LightCycler 96 detector (Roche). Expression levels were normalized to the housekeeping gene Beta-2 Microglobulin (B2M) using the comparative Ct-method 2^−ΔΔCt^. Experiments were carried out in triplicates, and RT-qPCR cycle threshold (Cq) data are depicted as mean values. The following mouse primers were used: Beta2m F-CGTGCGTGACATCAAAGAGA, R-CGCTGGTTGCCAATAGTGAT and mTGFβ1 F-CCTTCCTGCTCCTCATGG, R-CGCACACAGCAGTTCTTCTC. The following human primers were used: Beta2m F-ATGAGTATGCCTGGCCGTGTGA, R-GGCATCTTCAAACCTCCATG and hTGFβ1 F-TGTGCCCGGCTGCTGAAAGC, R-ACAGCAGCCCGAAGGGTCTCA.

### ELISA

To detect TGFβ1 released levels in conditioned media from all culture conditions we used a Human/Mouse TGFβ1 Quantikine ELISA kit (R&D systems). Culture conditioned media from all conditions were harvested after 5 days and frozen at -80°C until the assay. To ensure precision and reliability, we included a basal medium with BME as a control (Ctrl) to account for the potential presence of TGFβ1 in BME and its impact on T cell activation and Treg cell induction. We tested conditioned media from activated CD4^+^ T cells (CD4^+^) to specifically assess TGFβ1 secretion by these cells. Additionally, we included media from CRC tumor organoids cultured both with and without CD4^+^ T cells in our analysis. Latent TGFβ1 present in both controls and samples specimens was activated to immunoreactive TGFβ1 following the ELISA kit manufacturer’s protocol. Standards, controls and samples were then assayed in duplicate according to the manufacturer’s protocol. The optical density of the color was measured with a microplate reader set at 450 nm.

### Proliferation assays

For the assessment of murine CD4^+^ T cell proliferation, CD4^+^ T cells were stained with CellTrace Violet (CTV, Thermo Fisher) following the manufacturer’s protocol before initiating the culture. After 5-day incubation period detailed in the preceding culture section, CD4^+^ T cells were prepared for flow cytometry analysis. To evaluate the immunosuppressive effect of Treg cells on peripheral blood mononuclear cells (PBMCs) proliferation, CTV-stained PBMCs were co-cultured with TO-iTreg cells or TGFβ iTreg cells, harvasted after 5 days of culture, at different ratios (1:1, 1:2, 1:5 and 1:10). Experimental conditions included PBMCs alone (stained or unstained), CTV-stained PBMCs with TGFβ iTreg cells, CTV-stained PBMCs with TGFβ iTreg cells adjusted to match TI-Treg proportion, and CTV-stained PBMCs with TI-Treg cells. After 4 days of incubation, the cells were collected and subjected to flow cytometry analysis.

### Flow cytometry analysis

Murine and human CD4^+^ T cell were first stained with live/dead dye Zombie NIR (BioLegend) in PBS for 15 min at room RT, followed by subsequent staining with fluorochrome-labeled antibodies. Murine CD4^+^ T cells were stained with the following antibodies: anti-CD4-APC (BioLegend), anti-CD25-Pacific blue (BioLegend), anti-OX40-PE (eBioscience), anti-LAG3-PE (eBioscience), anti-CD27-PE-Cy7 (eBioscience) anti-GITR-PE-Cy7 (BioLegend), and anti-FAS-BV605 (BD optibuild) in MACs buffer for 15 min at RT. Human CD4^+^ T cells were stained with anti-CD4-FITC (BioLegend), anti-CD25-APC (BioLegend). This staining process occurred in MACs buffer for 15 minutes at room temperature, shielded from light. To evaluate human Treg cell differentiation, CD4^+^ T cells underwent intracellular staining using the Foxp3/Transcription Factor Staining Buffer Kit (Thermo Fisher Scientific) following the manufacturer’s protocol. Initially, cells were fixed for 45 minutes. Subsequently, anti-FOXP3-PE antibody (BD Biosciences), diluted in permeabilization buffer, was added and incubated for 30 minutes at room temperature, shielded from light. To characterize the markers expressed by hTO-iTreg cells, the same staining kit was utilized. However, FOXP3-PE-Cy7 antibody (eBioscience), diluted in permeabilization buffer, was applied overnight for intracellular staining. Flow cytometry data were acquired using a BD LSRFortessa Cell Analyzer (BD Biosciences) with FACSDiva (BD Biosciences) software and a CytoFLEX Flow Cytometer (Beckman Coulter) with CytExpert software. The data were analyzed with FlowJo (v.10.8.1, Treestar).

### RNA-sequencing and analysis

For RNA sequencing experiments, murine CD4^+^ T cell that were either cultured alone or with CRC tumor-organoids for five days, collected from the inserts and washed with pre-cold PBS. Zombie NIR, as a live/dead cell marker, and anti-CD4-APC were added to all single cell suspensions, as described before. Viable cells were FACS-sorted into 1.5ml eppendorfs based on CD4 and Foxp3 eGFP expression using the BD FACSAria II. Sorted cells were lysed in 100 μl TRIzol (Invitrogen) and stored at -80°C before shipping them on dry ice to Single Cell Discoveries B.V. (Utrecht, The Netherlands). RNA extraction was performed using the standard protocol. Bulk-cell RNA sequencing was conducted using a modified CELSeq protocol ^72,73^. In summary, purified RNA samples were uniquely barcoded with CEL-seq primers in a reverse transcription reaction ^72,73^. Then, barcoded cDNA molecules were pooled and linearly amplified through an *in vitro* transcription reaction. Amplified RNA was fragmented, and its quality assessed (Agilent bioanalyzer) before starting with the library preparation. Then, sequencing libraries were prepared with another reverse transcription reaction and PCR amplification to incorporate the right adapters for sequencing Illumina Truseq small RNA primers. The resulting sequenceable cDNA library underwent a quality assessment (Agilent bioanalyzer) prior to the paired-end sequencing in an Illumina Nextseq 500 System. The sequencing depth was of 10 million reads per sample. To commence the analysis, the Illumina library index and CEL-Seq sample barcode were initially identified. Subsequently, sequencing reads were aligned to the mm10 mouse transcriptome (RefSeq: NM_016701.3) employing the Burrows-Wheeler Alignment (BWA) tool ^74^. The mapping process and the generation of count tables were executed using the MapAndGo script available at https://github.com/anna-alemany/transcriptomics/tree/master/mapandgo. Counts were normalized for sequencing depth and RNA composition, using DESeq2’s median of ratios method^75^.

The RNA sequencing data were further analyzed using DESeq2 within the R2: Genomics Analysis and Visualization Platform (http://r2.amc.nl). This web application facilitated Principal Component Analysis (PCA, log2 z-score transformation), identification of differentially expressed genes (DEGs, p value ≤ 0.05), and the creation of heatmaps for specific gene sets displaying the mean z-score per sample. To determine significant changes in gene expression, the false discovery rate (FDR) method was applied for multiple testing correction, with a threshold of p value ≤ 0.05 (ANOVA or t-test) utilized to define significance. Venn diagrams, illustrating overlaps between DEG lists, were generated using the Bioinformatics & Evolutionary Genomics website (https://bioinformatics.psb.ugent.be/webtools/Venn/). Additionally, pathway analysis was conducted using DEGs as input in Toppfun (http://toppgene.cchmc.org) with pre-determined settings, focusing on gene ontology annotations for biological processes. Finally, to generate the progression-free interval (PFI) and overall survival (OS) curves, gene expression data of the TCGA cohort was filtered to include only genes upregulated TO-iTreg cells (top 141 genes) and then standardized using z-scores. Each patient’s mean z-score was calculated, and patients were divided into two equivalent groups (low and high expression) based on these scores. Patients with matching survival data were retained, and their group assignments were added to the survival data. A survival analysis was performed to compare outcomes between the low and high expression groups.

### Statistical analysis

Data analysis executed using GraphPad Prism 10. Mean values with corresponding standard deviations (SD) are presented, and a minimum of three experiments were conducted for each group (refer to Figure Legends for additional details). Statistical significance was determined through paired or unpaired one-way ANOVA with Tukey’s multiple comparison test or a student *t* test where appropriate. Significance levels are denoted by asterisks: *P* ≤ 0.05 (*), *P* ≤ 0.01 (**), or *P* ≤ 0.001 (***) or *P* < 0.0001 (****).

## Data availability statement

All data supporting the findings of this study are available from the lead contact upon reasonable request. RNA sequencing data analyzed in this manuscript have been deposited at GEO and are publicly available as of the date of publication.

