## supplementary figures legends for "Tumor-derived colorectal cancer organoids induce a unique Treg cell population through direct modulation of CD4^+^ T cell differentiation"

**Supplemetary Figures**

**Supplemental Figure 1. Flow cytometry analysis** **of Foxp3 eGFP^+^ Treg cells in transwell culture.**

(**a-b**) Representative flow cytometry gating strategy followed for assessing the percentage of CD4^+^ T cells expressing CD25 and Foxp3 eGFP in the conditions. The gating was performed under two conditions: (**a**) activated CD4^+^ T cells and (**b**) mTO-iTreg cells. (**c**-**d**) Percentage of CD4^+^ CD25^hi^ Foxp3 eGFP^+^ Treg cells among (**c**) activated CD4^+^ T cells or (**d**) TGFβ-iTreg cells cultured in the transwell insert alone (TW) or with matrix (BME) at the bottom of the plate, after 2 or 5 days of culture. Data represented as mean ± SD. P-values were calculated using one-way ANOVA. *p ≤ 0.05, **p ≤ 0.01, ***p ≤ 0.001 and ****p ≤ 0.0001, ns; not significant. CD4^+^: T activated cells. mTO1 and mTO2: CRC murine tumor-organoid lines.

**Supplemental Figure 2*.* Enhanced suppressive capacity of hTO-iTreg cells is lactate-independent.**

(**a-b**) Percentage of CD4^+^ CD25^hi^ FOXP3^+^ Treg cells after 5 days of CD4^+^ T cells culture alone or in co-culture with hTO1 and hTO2, supplemented with (**a**) sodium L-lactate (Lactate, 10mM) and (**b**) LHD inhibitor GSK2837808A (LDHi, 10µM), assessed using flow cytometry. (**c**) Expansion index and histograms showing the proliferation of CTV-stained PBMCs cultured alone or co-cultured with different ratios of diluted 30% TGFβ iTreg cells, 30% hTO1-iTreg cells, 30% hTO2-iTreg cells and 80% TGFβ iTreg cells after 4-day co-culture, assessed using flow cytometry (n = 3, different donors). Data represented as mean ± SD. P-values were calculated using one-way ANOVA. *p ≤ 0.05, **p ≤ 0.01, ***p ≤ 0.001 and ****p ≤ 0.0001, ns; not significant. CD4^+^: T activated cells, hTO1 and hTO2: CRC human tumor-organoid lines.

**Supplemental Figure 3*.* Immune regulation pathways shared by *in vitro* CRC TO-iTreg cells and TGFβ-iTreg cells.**

(**a-b**) Heatmaps of the 126 differentially (**a**) upregulated or (**b**) downregulated genes in TGFβ-iTreg cells (green) and mTO2-iTreg cells (red) compared to eGFP^-^ CD4^+^ T cells (blue) (ANOVA, FDR, p ≤ 0.05). (**c**) GO term analysis focus on biological processes associated with the 126 genes shared between TGFβ-iTreg cells and mTO2-iTreg cells, as compared to CD4^+^ eGFP^-^ T cells (ANOVA, FDR, p ≤ 0.05). The numbers inside the bars represent the number of genes represented in that specific GO term. P-values were calculated using one-way ANOVA. *p ≤ 0.05, **p ≤ 0.01, ***p ≤ 0.001 and ****p ≤ 0.0001, ns; not significant. mTO2: CRC murine tumor-organoid line.

**Supplemental Figure 4*.* Metabolic and signaling pathway differences in *in vitro* CRC TO-iTreg cell compared to TGFβ-iTreg cells.**

(**a**) Heatmap of the 56 most significantly DEGs between mTO2-iTreg cells compared to TGFβ-iTreg cells (ANOVA, FDR, p ≤ 0.01). (**b**) GO term analysis focus on biological processes associated with the 1155 DEGs between mTO2-iTreg cells and TGFβ-iTreg cells (ANOVA, FDR, p ≤ 0.05). The numbers inside the bars represent the number of genes represented in that specific GO term. (**c**) Heatmap and (**d**) bar chart quantifying (mean ± SD) the signature z-score of the DEGs associated with the GO glycolysis in mTO2-iTreg cells compared to TGFβ-iTreg cells. (**c**) Heatmap and (**d**) bar chart quantifying (mean ± SD) the signature z-score of the DEGs associated with the GO glycolysis in mTO2-iTreg cells compared to TGFβ-iTreg cells. (**e**) Bar chart quantifying (mean ± SD) the signature z-score of the expression of TFs genes associated with Tregs cells in mTO2-iTreg cells compared to TGFβ-iTreg cells. (**f**) Bar chart quantifying (mean ± SD) the signature z-score and heatmap of the expression of genes associated with *in vivo* CRC TI-Treg cells in mTO2-iTreg cells compared to TGFβ-iTreg cells. P-values were calculated using one-way ANOVA and paired t-test analysis. *p ≤ 0.05, **p ≤ 0.01, ***p ≤ 0.001 and ****p ≤ 0.0001, ns; not significant. mTO2: CRC murine tumor-organoid line.

**Supplemental Figure 5*.* 141 upregulated genes in mTO2-iTreg cells with prognostic value.**

Heatmap of the 141 most upregulated genes (with >3 log-fold change) in mTO2-iTreg cells compared to TGFβ-iTreg cells used for the prognostic curves (ANOVA, FDR, p ≤ 0.05). P-values were calculated using one-way ANOVA. mTO2: CRC murine tumor-organoid line.
