## supplementary figures for "Tumor-derived colorectal cancer organoids induce a unique Treg cell population through direct modulation of CD4^+^ T cell differentiation"

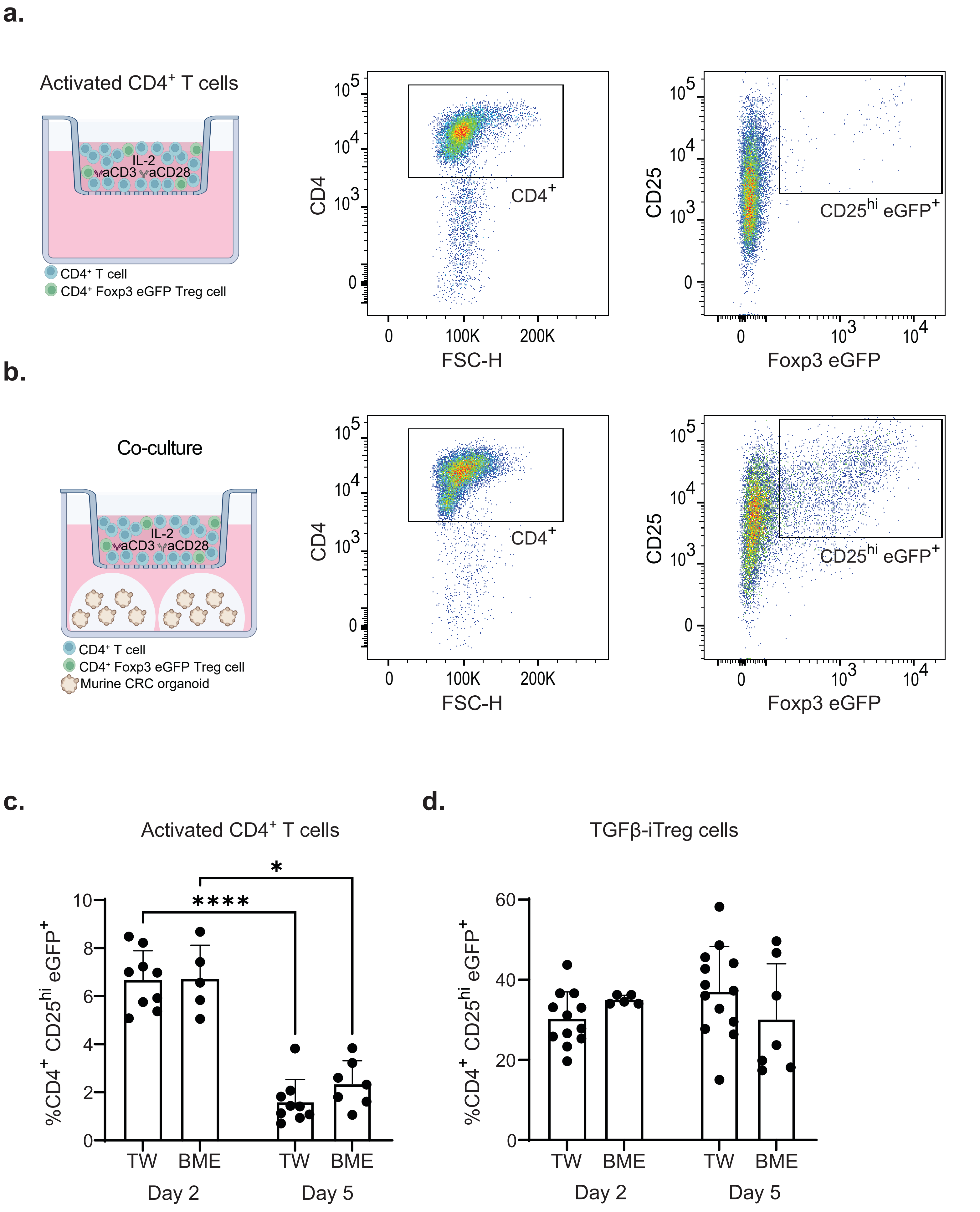
a.
Activated CD4+ T cells
IL-2
aCD3
aCD28
CD4+ T cell
CD4+ Foxp3 eGFP Treg cell
CD4
FSC-H
CD4+
CD25
Foxp3 eGFP
CD25hi eGFP+
b.
Co-culture
IL-2
aCD3
aCD28
CD4+ T cell
CD4+ Foxp3 eGFP Treg cell
Murine CRC organoid
CD4
FSC-H
CD4+
CD25
Foxp3 eGFP
CD25hi eGFP+
c.
Activated CD4+ T cells
%CD4+ CD25hi eGFP+
TW
BME
TW
BME
Day 2
Day 5
d.
TGFβ-iTreg cells
%CD4+ CD25hi eGFP+
TW
BME
TW
BME
Day 2
Day 5

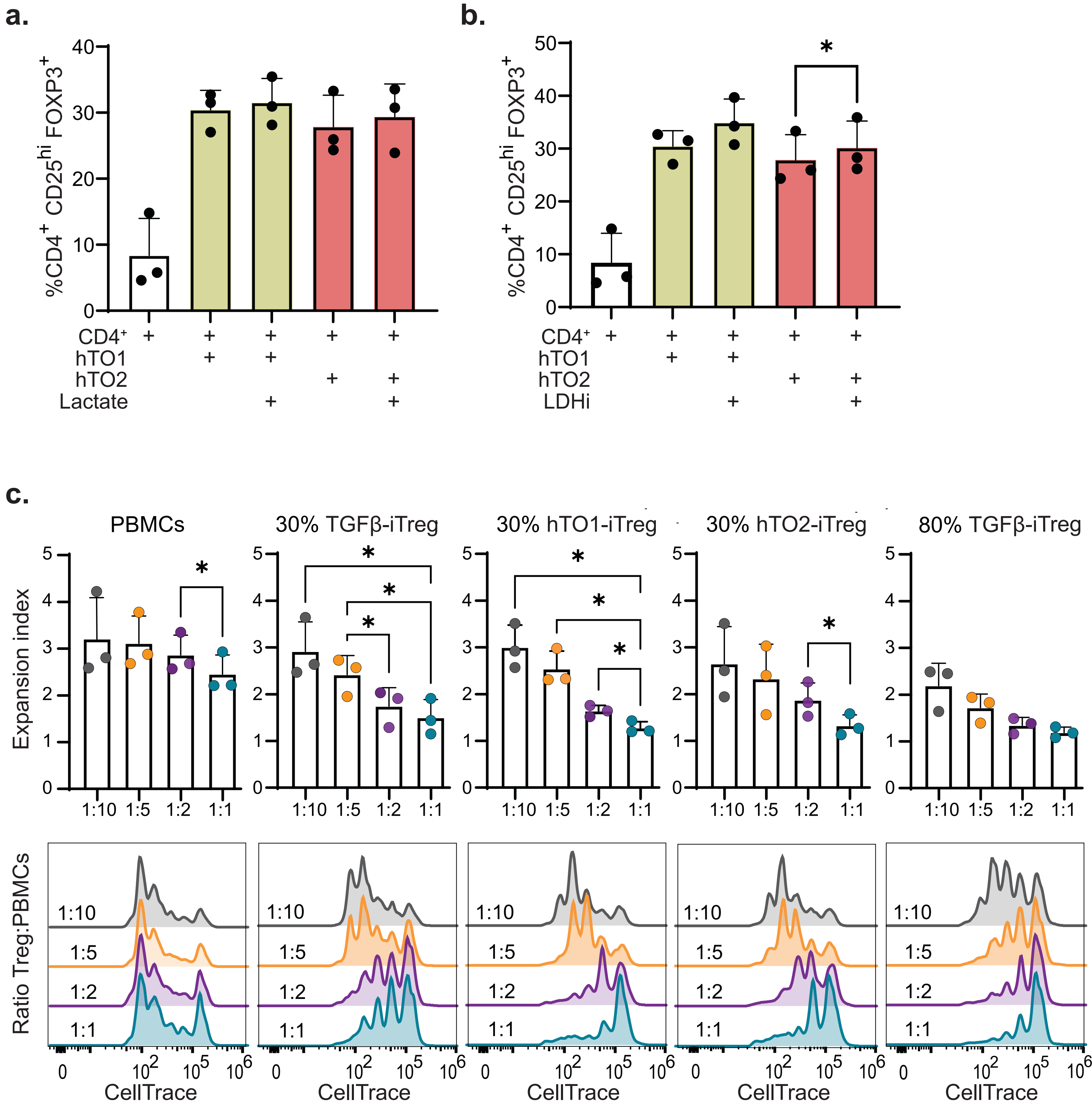


Supplementary Figure 2

Aristin Revilla S. *et al.*

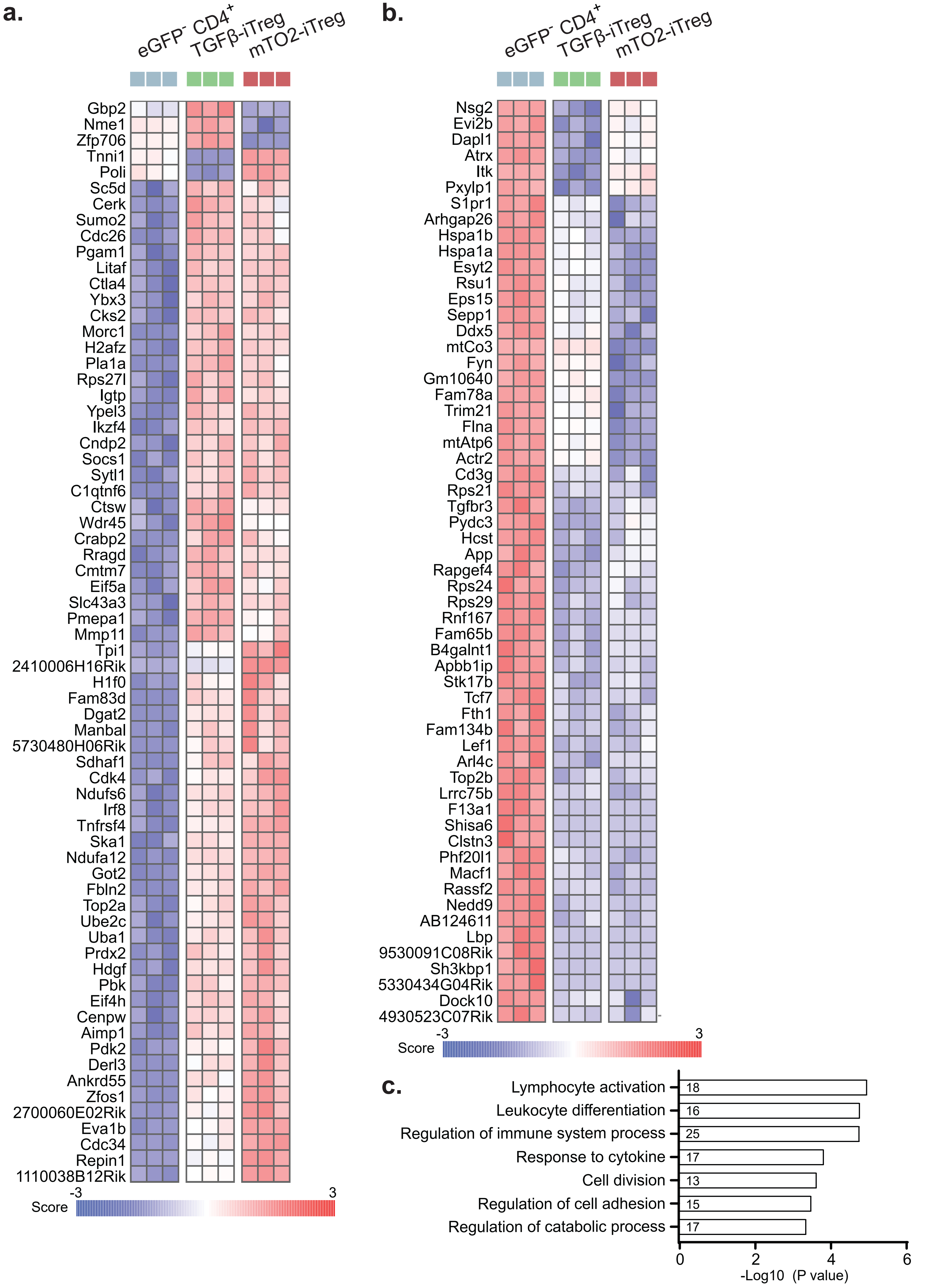


Supplementary Figure 3

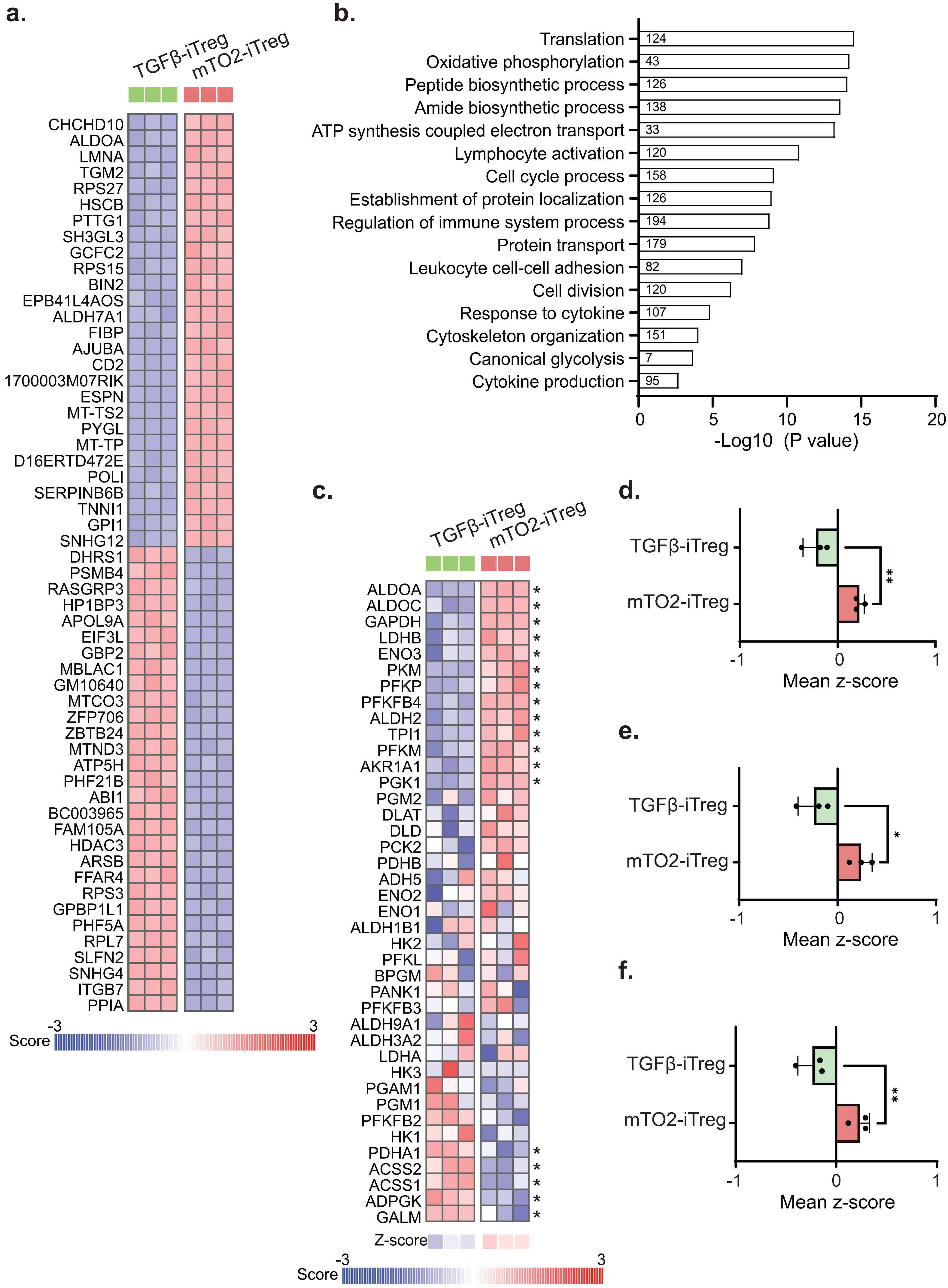

a.
TGFβ-iTreg
mTO2-iTreg
CHCHD10
ALDOA
LMNA
TGM2
RPS27
HSCB
PTTG1
SH3GL3
GCFC2
RPS15
BIN2
EPB41L4AOS
ALDH7A1
FIBP
AJUBA
CD2
1700003M07RIK
ESPN
MT-TS2
PYGL
MT-TP
D16ERTD472E
POLI
SERPINB6B
TNNI1
GPI1
SNHG12
DHRS1
PSMB4
RASGRP3
HP1BP3
APOL9A
EIF3L
GBP2
MBLAC1
GM10640
MTCO3
ZFP706
ZBTB24
MTND3
ATP5H
PHF21B
ABI1
BC003965
FAM105A
HDAC3
ARSB
FFAR4
RPS3
GPBP1L1
PHF5A
RPL7
SLFN2
SNHG4
ITGB7
PPIA
Score
-3
3
b.
Translation 124
Oxidative phosphorylation 43
Peptide biosynthetic process 126
Amide biosynthetic process 138
ATP synthesis coupled electron transport 33
Lymphocyte activation 120
Cell cycle process 158
Establishment of protein localization 126
Regulation of immune system process 194
Protein transport 179
Leukocyte cell-cell adhesion 82
Cell division 120
Response to cytokine 107
Cytoskeleton organization 151
Canonical glycolysis 7
Cytokine production 95
0 5 10 15 20
-Log10 (P value)
c.
TGFβ-iTreg
mTO2-iTreg
ALDOA *
ALDOC *
GAPDH *
LDHB *
ENO3 *
PKM *
PFKP *
PFKFB4 *
ALDH2 *
TPI1 *
PFKM *
AKR1A1 *
PGK1 *
PGM2
DLAT
DLD
PCK2
PDHB
ADH5
ENO2
ENO1
ALDH1B1
HK2
PFKL
BPGM
PANK1
PFKFB3
ALDH9A1
ALDH3A2
LDHA
HK3
PGAM1
PGM1
PFKFB2
HK1
PDHA1 *
ACSS2 *
ACSS1 *
ADPGK *
GALM *
Z-score
Score
-3
3
d.
TGFβ-iTreg
mTO2-iTreg
**
-1 0 1
Mean z-score
e.
TGFβ-iTreg
mTO2-iTreg
*
-1 0 1
Mean z-score
f.
TGFβ-iTreg
mTO2-iTreg
**
-1 0 1
Mean z-score

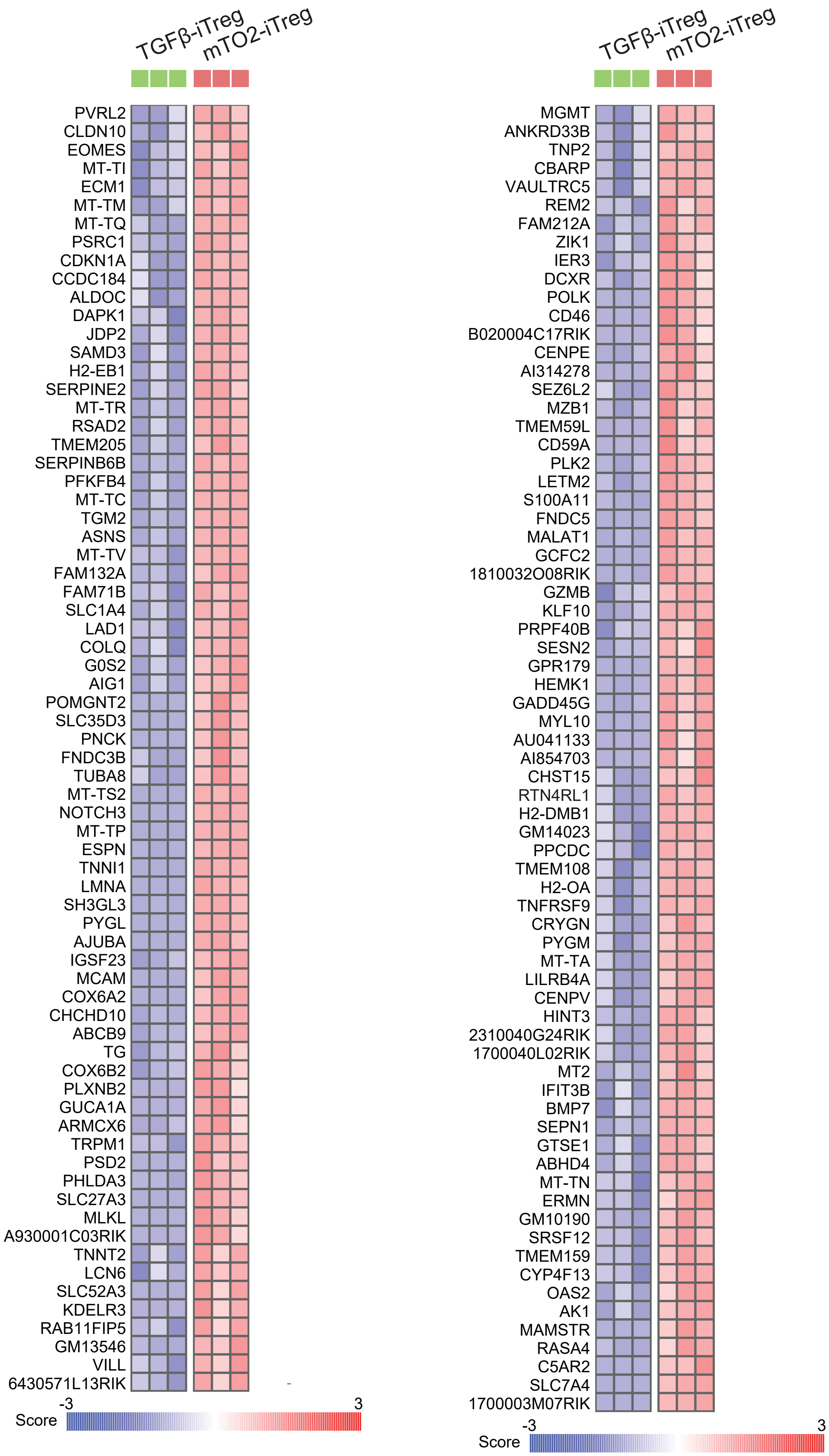


Supplementary Figure 5

Aristin Revilla S. *et al.*
